# Reduced virulence and enhanced host adaption during antibiotics therapy: A story of a within-host carbapenem-resistant *Klebsiella pneumoniae* sequence type 11 evolution in a fatal scrotal abscess patient

**DOI:** 10.1101/2021.05.06.443049

**Authors:** Meiping Ye, Chunjie Liao, Mengya Shang, Danyang Zou, Jingmin Yan, Zhixiang Hu, Xiaogang Xu, Jianping Jiang, Pingyu Zhou

**Affiliations:** STD Institute, Shanghai Skin Disease Hospital, Tongji University School of Medicine, Shanghai, China; Shanghai Skin Disease Hospital, Clinical School of Anhui Medical University, Shanghai, China; Institute of Antibiotics, Huashan Hospital, Fudan University, Shanghai 200040, China

**Keywords:** Genomic study, *in vivo* evolution, host adaptation, RNA-Seq

## Abstract

Carbapenem-resistant *Klebsiella pneumoniae* (CRKP) has disseminated globally and become a major threat to human life. The sequence type (ST) 11 CRKP is a dominant clone in Asia, especially China, but how this clone evolves *in vivo*, then adapts to host and facilitates dissemination remain largely unknown. We analyzed the genomic dynamics of 4 ST11-CRKP isolates sequencially isolated from the urine of a patient with initial fatal scrotal abscess and finally recovered without effective medication. Genomic differences were identified and their implications for pathogenesis and host adaptation were investigated. The related transcriptional pathways were further explored by RNA-Seq. Genomic analysis identified 4-24 mutations and 94%-100% were synonymous or intergenic. The mutation rate of ST11-CRKP was 2.1×10^−6^-1.7×10^−5^ substitutions/site/year over 47 days of antibiotics therapy. During this period, CRKP underwent several adaptive changes including tigecycline resistance and virulence attenuation. Tigecycline resistance was caused by *ramR* ribosomal binding site (RBS) deletion, which has been described by us previously. In this study, we demonstrated that mutations associated with acyltransferase (*act*) and *ompK26* caused the virulence attenuation of ST11-CRKP*. act* deletion reduced the production of capsular polysaccharide and enhanced biofilm formation. RNA-Seq analysis revealed that *act* influenced the expression of *ldhA*, *bglX*, *mtnK* and *metE* which likely participate in capsular synthesis and biofilm formation. *ompK26* affected the virulence by its overexpression caused by the deletion of upstream repressor binding site. Our finding suggested that the broad genomic diversity, high evolutionary capacity and rapid within-host adaptability of ST11-CRKP might contribute to the worldwide dissemination of this clone.

**IMPORTANCE:** Carbapenem-resistant *Klebsiella pneumoniae* (CRKP) has disseminated worldwide and can cause life threatening infections, including pneumonia, bloodstream infections, urinary tract infections, intra-abdominal infection, liver abscess and meningitis. CRKP infection is the leading cause of high mortality in clinical. The sequence type (ST) 11 CRKP is a dominant clone and accounts for 60% of CRKP infections in China. Recently, the ST11-CRKP with high transmissibility are increasingly identified. Understanding how this clone evolved is crucial in controlling its further dissemination. The significance of our research is identifying the *in vivo* genomic dynamics of ST11-CRKP and the genetic basis for ST11-CRKP to facilitate persistence and dissemination, which will has broader biomedical impacts on understanding of ST11-CRKP dissemination. Furthermore, our study also highlights the importance of monitoring the development of variation in antibiotics susceptibility and virulence of bacteria in clinical practice, considering that pathogens can rapidly adapt to host during the treatment.

## INTRODUCTION

The worldwide dissemination of carbapenem-resistant *Enterobacteriaceae* (CRE) has become an urgent threat to public health (1) and CRE has been classified as critical on the global priority pathogens list by the World Health Organization (2). Carbapenem-resistant *Klebsiella pneumoniae* (CRKP) is the most common genus of CRE. Study showed that CRKP infection accounts for 80% of CRE infection and is the leading cause of high mortality in clinical infections (3).

Most of the CRKP belong to the clonal group CG258, in which sequence type (ST) 258 and ST11 are dominant (4). *K. pneumoniae* ST11 is the predominant clone in Asia and accounts for 60% of CRKP in China (5). Recently, a subclone of ST11-CRKP with high transmissibility is increasingly identified in China (6). Though with the extensive interrogations, the evolutionary success of ST11-CRKP for dissemination is not fully understood. The known factors impacting the dissemination includes resistance to carbapenem, high transmissibility, pathogenicity alternation and increased duration (7, 8). Besides, host adaptation or adaptibility played a crucial role in clone transmission, which was highlighted by a study on the long-term carriage of ST258-CRKP within a patient (9). However, little is known about the within-host adaptation of ST11-CRKP in genomic and transcriptomic scales.

In this study, we analyzed the within-host evolution of ST11-CRKP isolated from the urine of a patient with initial fatal scrotal abscess and finally recovered without effective medication. During this course, the ST11-CRKP incurred a series of phenotypic variations including tigecycline resistance, virulence attenuation, capsular polysaccharide (CPS) reduction and biofilm formation enhancement, and then its adaptation to the host environment was increased. However, genomic analysis revealed that the strain underwent limit genetic changes. The deletion of *ramR* ribosomal binding site (RBS) mediated the tigecycline resistance, which has been described in our previous study (10). Therefore, here we further characterized other genomic changes associated with phenotypic variations and interrogated them using wet-lab experiments, transcriptome analysis and animal infection models. Our results make a connection between genomic variation and within-host adaption, which will help deepen the knowledge and understanding of ST11-CRKP dissemination. In addition, results from our study also highlights the importance of monitoring the development of variation in antibiotics susceptibility and virulence of bacteria in clinical practice, considering that pathogens can rapidly adapt to host during the treatment.

## RESULTS

### The virulence of CRKP was attenuated in vivo during the antibiotics therapy

The four CRKP isolates (KP-1S, KP-2S, KP-3R and KP-4R) were sequentially isolated from the urine of a 50-year-old male patient with scrotal abscess during 47 days of tigecycline-containing antibiotics therapy. The initial isolates (KP-1S and KP-2S) were susceptible to tigecycline and subsequent isolates (KP-3R and KP-4R) were resistant to tigecycline. All the four CRKP isolates were resistant to 17 other antimicrobials including amikacin, gentamicin, ciprofloxacin, norfloxacin, sulfamethoxazole, nitrofurantoin, piperacillin, TXP, cefazolin, cefuroxime, cefotaxime, ceftazidime, cefepime, cefoperazone/ sulbactam, cefmetazole, imipenem and meropenem except polymyxin B. At that time, polymyxin B was not approved by National Medical Products Administration in China.

As the condition of the patient did not get worse after the isolation of KP-4R, we concluded that the virulence of the subsequent isolates had reduced. To test our hypothesis, we performed an animal experiment with the four isolates. As showed in **Figure 1**, the mortality rate of mice inoculated with KP-1S (90%) or KP-2S (90%) was significantly higher (P < 0.05) than that of mice inoculated with KP-3R (20%) or KP-4R (30%), demonstrating that the virulence of KP-3R and KP-4R had reduced.

**Figure 1.**
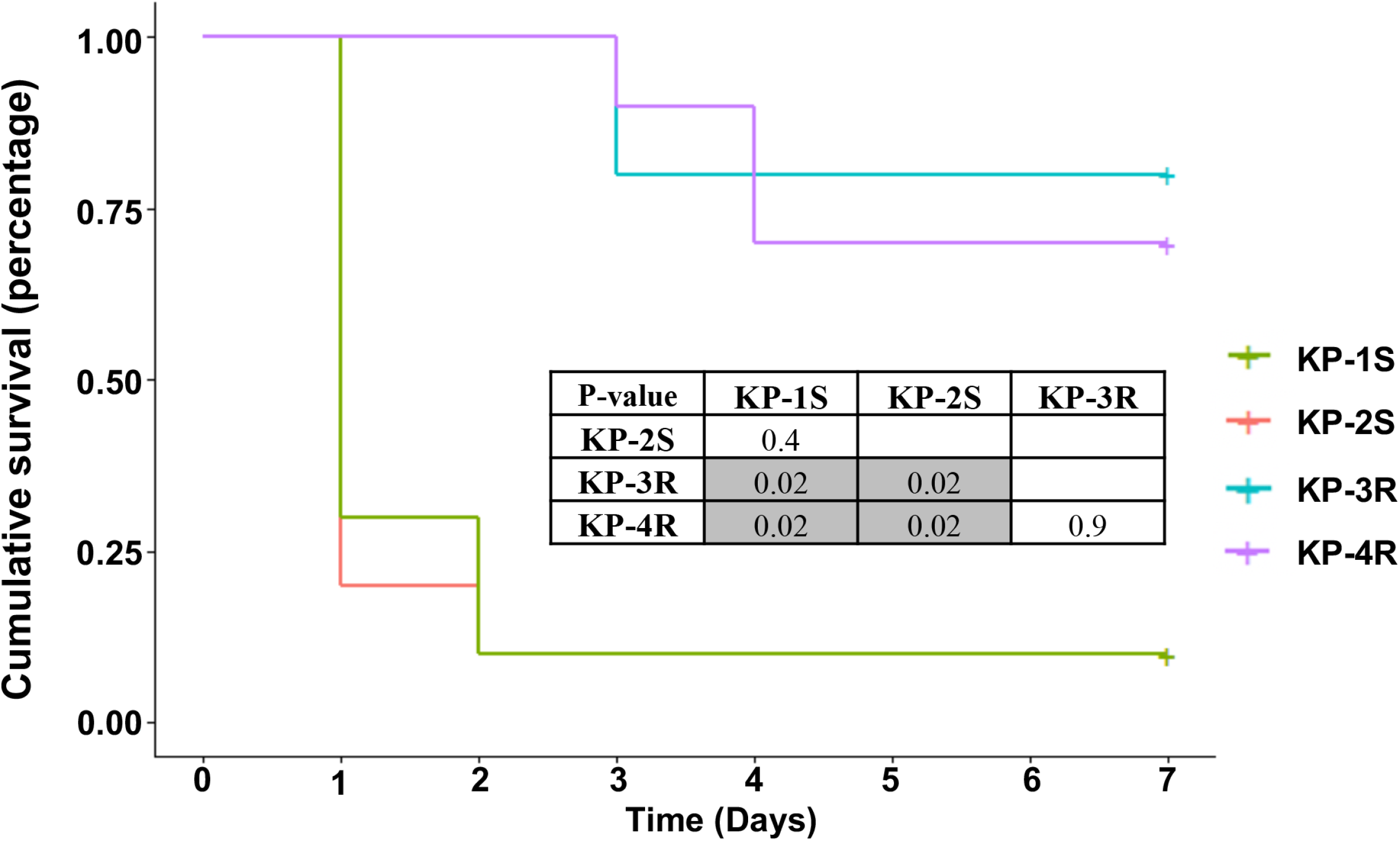
Kaplan-Meier survival curves of mice intraperitoneally challenged with *Klebsiella pneumoniae* strains KP-1S, KP-2S, KP-3R and KP-4R. Ten mice in each group inoculated with 10^6^ CFUs were monitored daily for 7 days. P values were calculated from the Mantel-Cox log rank test for survival curve comparison. Grey shading indicates significant values (<0.05).

### CRKP isolates harbor virulence genes including *aerobactin*, *rmpA* and *rmpA2*

To investigate the genomic features of the four CRKP isolates, we sequenced the whole genomes of KP-1S, KP-2S, KP-3R and KP-4R. Genomic analysis showed that all the CRKP isolates belong to the KL64-ST11, and all of them harbor carbapenemase-encoding gene *bla*_KPC-2_ and extended-spectrum β-lactamase-encoding gene *bla*_CTX-M-65_, which confer resistance to carbapenem and cephalosporin. Furthermore, besides yersiniabactin which is usually located on the chromosome of *K. penumoniae*, *aerobactin* (*iutAiucABCD*), *rmpA* and *rmpA2* that are mostly identified on a large virulence plasmid (11) were found on the genomes of all the CRKP isolates (**Table 1**), indicating that the virulence of CRKP in this study has been enhanced compared to classic ST11-CRKP. Plasmid replicon analysis was identified as ColRNAI, IncFIB(K), IncFII, IncHI1B and IncR in each CRKP isolates. Among them, IncFIB(K) and IncHI1B were usually associated with the hypervirulent plasmid in hypervirulent *K. penumoniae*, while IncFII and IncR are usually associated with *bla*_KPC-2_.

**Table 1.** Genomic features of KP-1S, KP-2S, KP-3R and KP-4R

| Isolates | MLST | KPC | ESBL | wzi | Yersiniabactin | Aerobactin | rmpA | rmpA2 | Plasmid Inc |
| --- | --- | --- | --- | --- | --- | --- | --- | --- | --- |
| KP-1S, KP-2S,<br>KP-3R and<br>KP-4R | ST11 | KPC-2 | CTX-M-65 | wzi64 | ybt 9<br>(ICEKp3) | iuc 1 | rmpA-2<br>(KpVP-1) | rmpA2-3<br>(-47%) | ColRNAI, IncFIB(K), IncFII, IncHI1B,<br>IncR |

### ST11-CRKP strain underwent limit genetic changes during the period in the host

Whole genome alignments showed that no genomic rearrangement has happened to KP-2S, KP-3R or KP-4R (**Figure 2A**) compared with KP-1S. We further identified the variants including SNPs and INDELs among the four CRKP isolates using KP-1S as the reference (**Figure 2B**). In KP-2S, 1 SNP and 3 INDELs were found. Among them, the SNP is a synonymous variant and all the 3 INDELS are located in the intergenic regions. In KP-3R, 11 SNPs and 13 INDELS were found. Among them, 3 SNPs are synonymous variants and 8 are located in the intergenic regions. Except for the deletion containing acyltransferase (*act*) family protein, the other 11 INDELs are located in the intergenic regions. In KP-4R, 10 SNPs and 9 INDELs were found. Among them, 1 SNP is a synonymous variant and 9 are located in the intergenic regions. Like KP-3R, all the INDELs are located in the intergenic regions except for the deletion containing *act*. The instantaneous mutation rates for KP-2S, KP-3R and KP-4R are 2.1×10^−6^ substitutions per site per year, 1.7×10^−5^ substitutions per site per year and 1.3×10^−5^ substitutions per site per year, respectively (**Figure 2C**).

**Figure 2.**
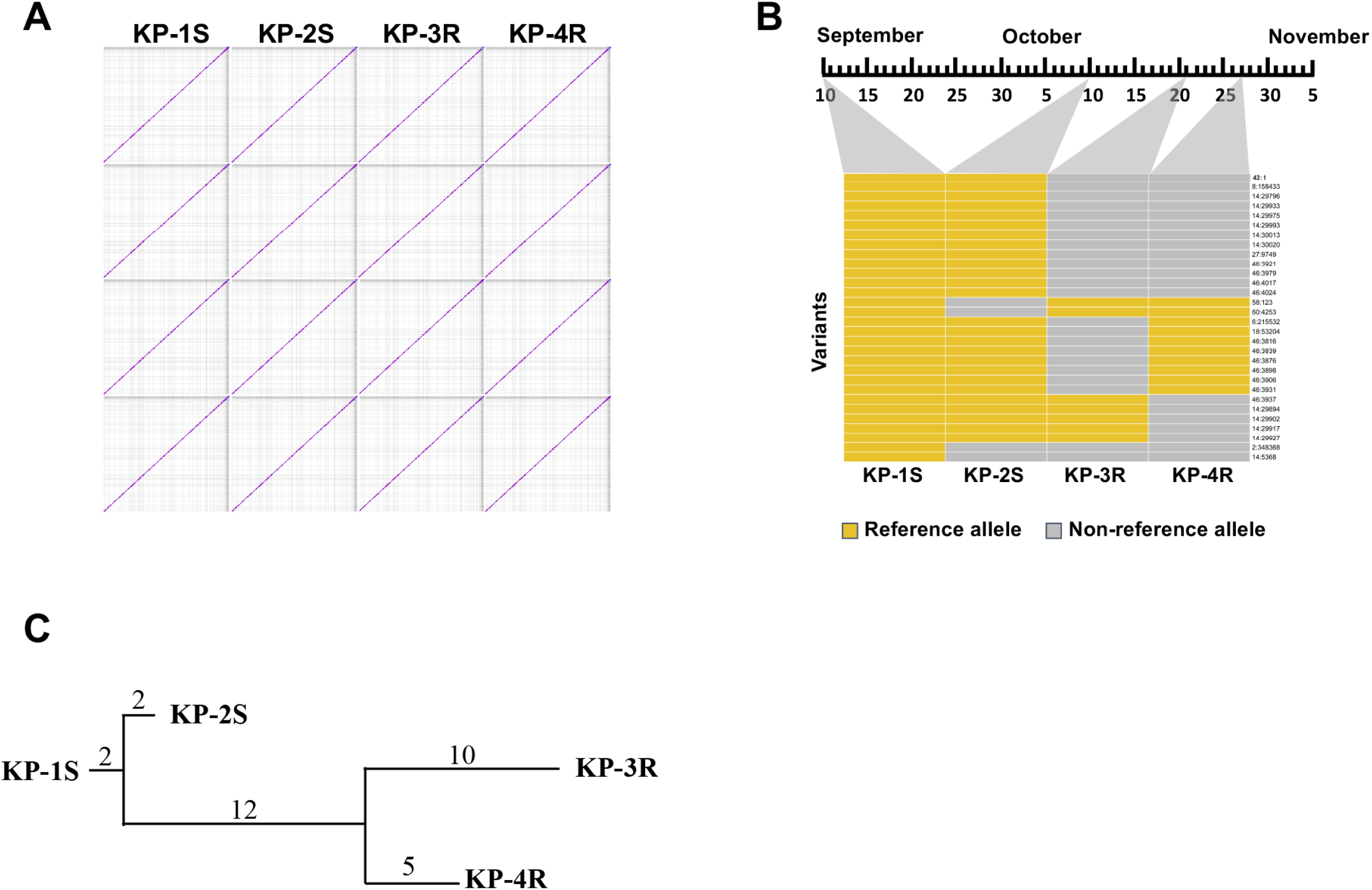
Genomic analysis of KP-1S, KP-2S, KP-3R and KP-4R. A. The pairwise genome alignments of KP-1S, KP-2S, KP-3R and KP-4R. B. Genomic variants of each CRKP isolates. C. The phylogenetic tree of the four CRKP isolates based on the genomic variants. The numbers above the line represented the number of mutations. Mutation rates were calculated according to the number of variations and isolation time-span.

To explore the genetic determinants associated with the phenotypic changes, variants present in KP-3R and KP-4R, and absent in KP-1S and KP-2S were further considered, and shown in **Table 2**. A total of 14 variants were exclusively present in KP-3R and KP-4R, including 8 SNPs and 6 INDELs. All of the variants are located in the intergenic regions, except for the 2,226 bp large deletion which contains *act*. The 2,226 bp large deletion is located upstream of an insertion sequence *ISKpn26* which usually mediates DNA inversion or deletion in *K. pneumoniae* (12) (**Figure S1**). Besides the 2,226 bp large deletion, the 12 bp-deletion of *ramR* RBS and the 5 bp (TGTTT)-deletion 42 bp upstream of *ompK26*, other 11 variants are located either on the downstream of or far away (> 200bp) from their adjacent genes and thus were considered not essential for phenotypic changes. We previously demonstrated that the 12 bp-deletion of *ramR* confers tigecycline resistance (10), and studies revealed that the impact of *ramR* on pathogenicity is limited (13, 14). Therefore, in this study, we focused on the functions of the TGTTT deletion and the *act* deletion, which likely affects the virulence of KP-3R and KP-4R.

**Table 2.**
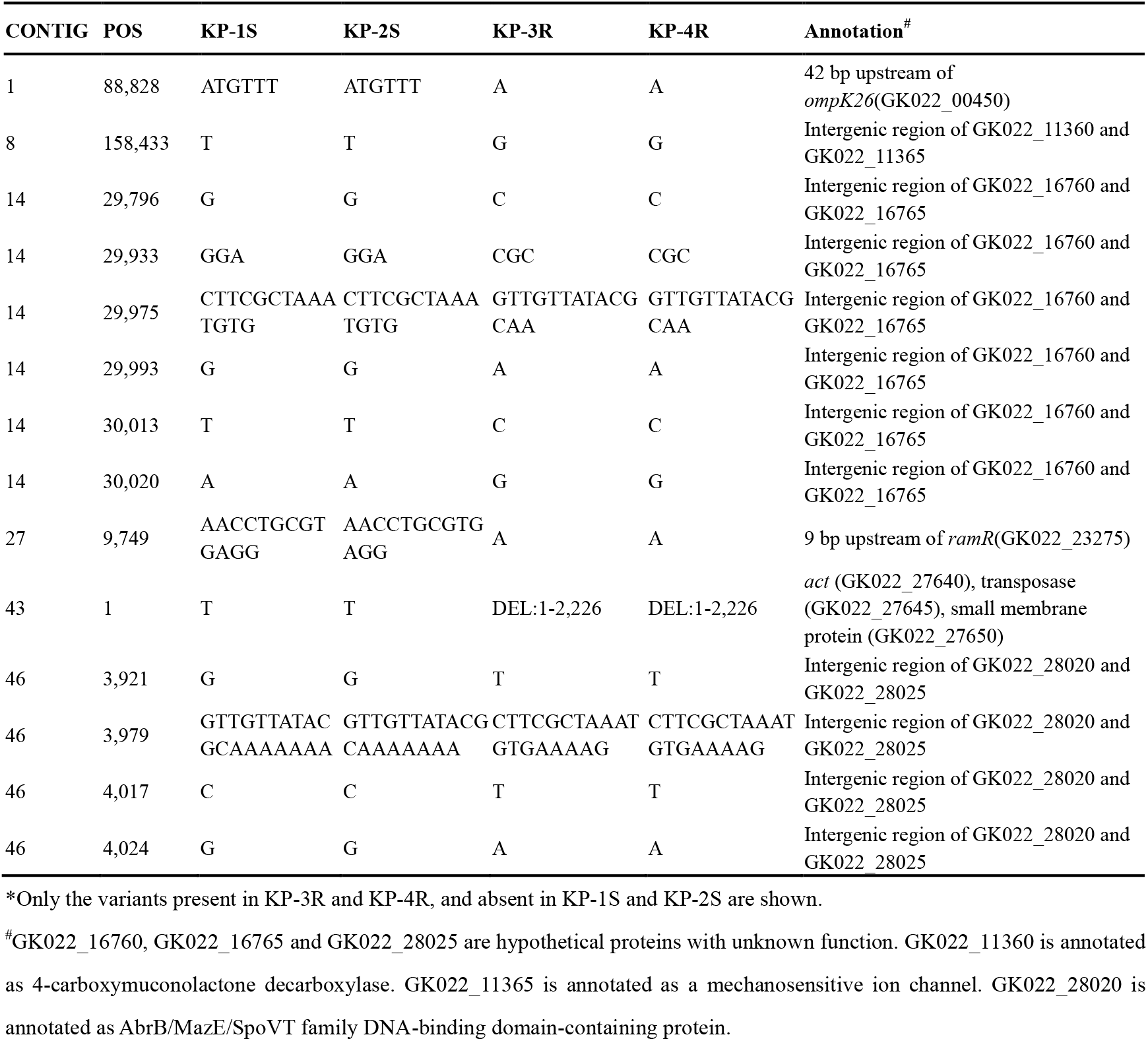
Genomic differences among the four isolates*

### TGTTT deletion upregulated the expression of *ompK26* by destroying the binding site of repressor KdgR and partially reduced the virulence of CRKP

qRT-PCR showed that *ompK26* was significantly over-expressed in KP-3R and KP-4R compared with KP-1S and KP-2S (**Figure 3A**). To validate whether the *ompK26* over-expression was caused by TGTTT deletion, *ompK26* with its native and mutant promoter regions were cloned in a T-vector to generate pMY53 and pMY54 (**Figure S2**) and transformed into KP-3R△*ompK26* (**Figure S3**), respectively. As shown in **Figure 3B**, the transcriptional level of *ompK26* in KP-3R△*ompK26*/pMY54 was significantly higher than that in KP-3R△*ompK26*/pMY53, demonstrating that the deletion of TGTTT upregulated the *ompK26* expression.

**Figure 3.**
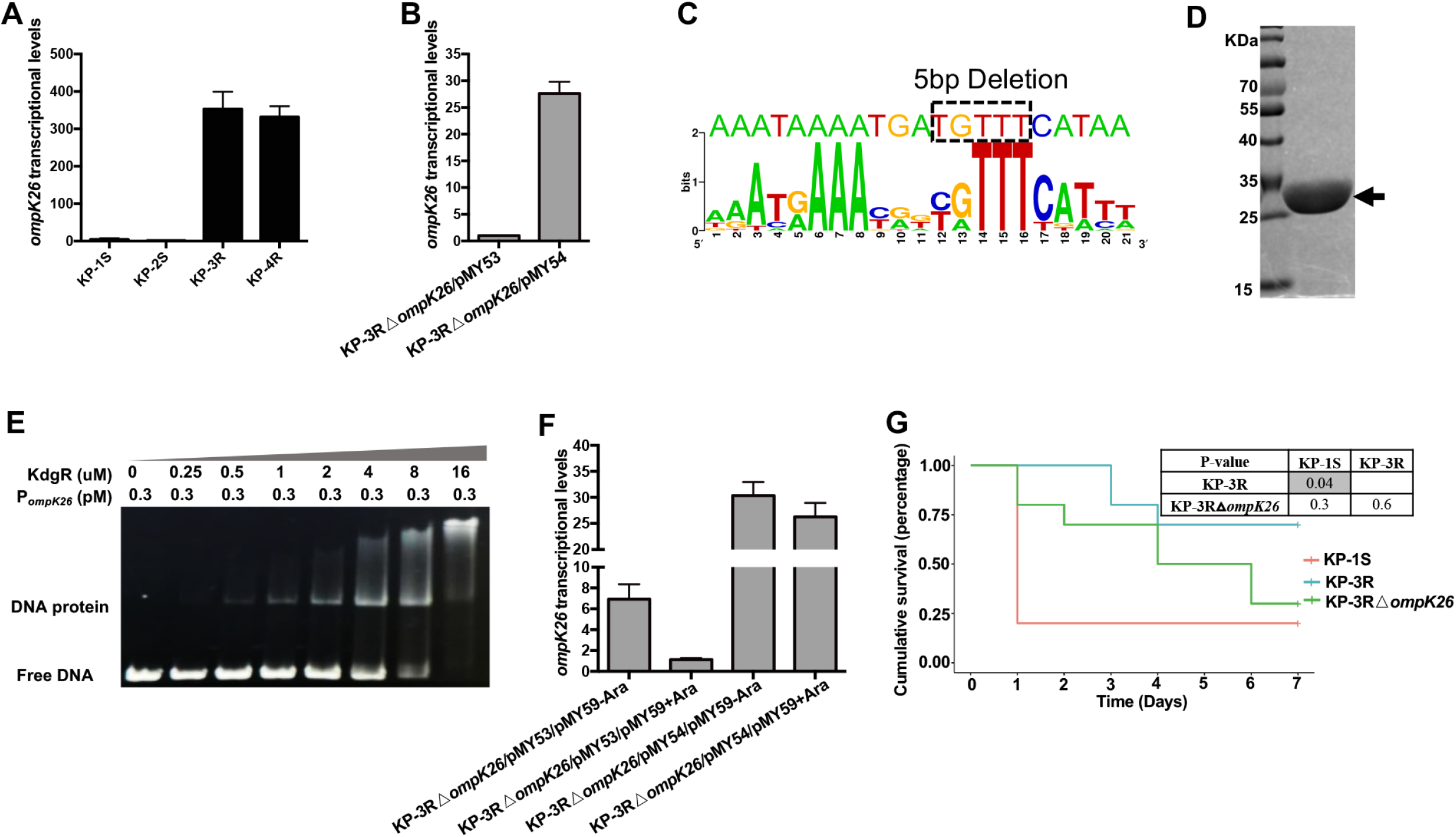
The functional study of *ompK26.* A. Quantitative reverse transcription PCR (qRT-PCR) assessment of the transcriptional level of *ompK26* in KP-1S, KP-2S, KP-3R and KP-4R. B. qRT-PCR assessment of the transcriptional level of *ompK26* in KP-3R△*ompK26*/pMY53 and KP-3R△*ompK26*/pMY54. C. The prediction of KdgR binding site. D. KdgR-6XHis protein following affinity purification. The arrow indicates KdgR protein. E. EMSA using purified KdgR protein. KdgR decreases the migration of promotor DNA of *ompK26*. F. qRT-PCR assessment of the transcriptional level of *ompK26* in KP-3R△*ompK26*/pMY53/pMY59-Ara, KP-3R△*ompK26*/pMY53/pMY59+Ara, KP-3R△*ompK26*/pMY54/pMY59-Ara and KP-3R△*ompK26*/pMY54/pMY59+Ara. G. Kaplan-Meier survival curves of mice intraperitoneally challenged with KP-1S, KP-3R and KP-3R△*ompK26*. Mice were injected with 10^6^ CFUs and monitored for 7 days. P values were calculated from the Mantel-Cox log rank test for survival curve comparison. Grey shading indicates significant values (<0.05).

Given that OmpK26 belong to KdgM family, which is usually under the control of KdgR. Thus, the putative KdgR binding site upstream of *ompK26* was predicted. Result showed that TGTTT fell into the KdgR binding region **(Figure 3C**). To validate the prediction, KdgR was purified (**Figure 3D**) for EMSA experiment. The results showed that KdgR can bind to the promoter region of *ompK26* (**Figure 3E**). To further demonstrate that *ompK26* was under the regulation of KdgR *in vivo*, KP-3R△*ompK26*/pMY53 and KP-3R△*ompK26*/pMY54 were complemented with a wild-type KdgR (pMY59) under the control of the arabinose-inducible promoter P_BAD_. As shown in **Figure 3F**, when KdgR was induced in the presence of arabinose, the transcription of *ompK26* in KP-3R△*ompK26*/pMY53 was significantly repressed. However, the repressive effect was not observed in KP-3R△*ompK26*/pMY54. These results together demonstrated that TGTTT fell into the binding region of KdgR, and the deletion of TGTTT upregulated the expression of *ompK26*.

To investigate the role of OmpK26 in virulence, a mouse lethality study of KP-1S, KP-3R and KP-3R△*ompK26* was conducted. Results showed that, though without significance, the mortality rate of KP-3R△*ompK26* (70%) is between that of KP-1S (80%) and KP-3R (30%) (**Figure 3G**). These results indicated that *ompK26* was associated with virulence, and overexpression of *ompK26* slightly reduced the virulence of CRKP.

### *act* is involved in the synthesis of CPS and deletion of *act* significantly attenuated the virulence of ST11-CRKP

To explore the function of *act*, the *act* mutant strain KP-1S△*act* was constructed (**Figure S3**) and subjected to a mouse lethality test. As shown in **Figure 4A**, the mortality rate of mice infected with KP-1S△*act* (20%) was significantly lower (P < 0.05) than that of mice infected with KP-1S (80%). No significant difference in survival rate was found between the groups of KP-1S△*act* and KP-3R, indicating that *act* is an important virulence factor in ST11-CRKP.

**Figure 4.**
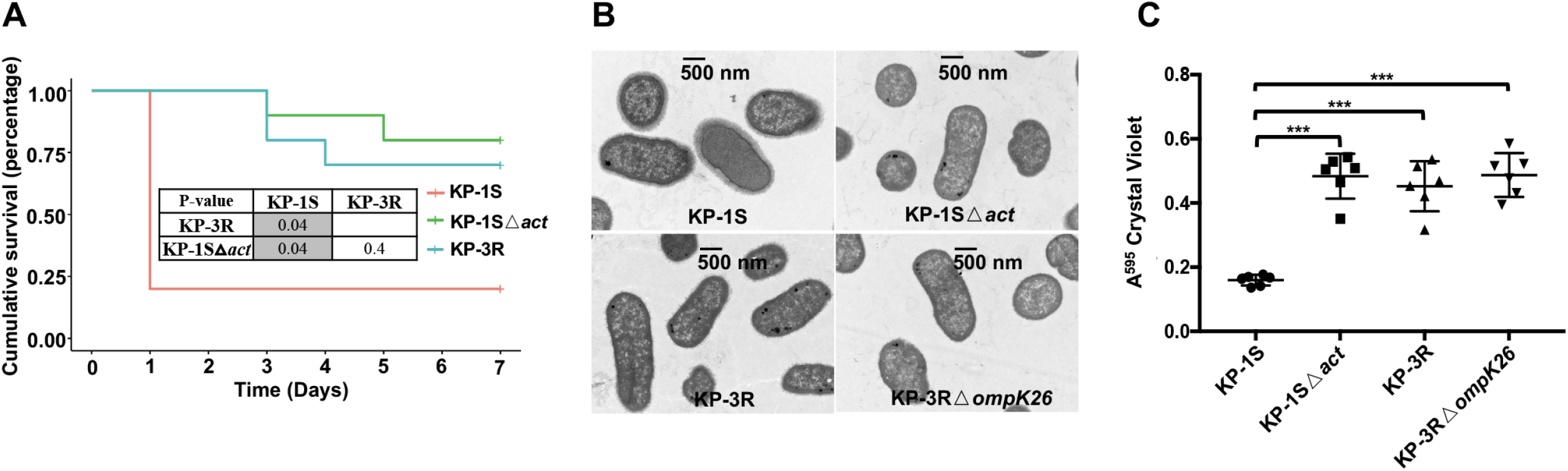
The functional study of *act*. A. Kaplan-Meier survival curves of mice intraperitoneally challenged with KP-1S, KP-1S△*act* and KP-3R. Mice were injected with 10^6^ CFUs and monitored for 7 days. P values were calculated from the Mantel-Cox log-rank test for survival curve comparison. Grey shading indicates significant values (<0.05). B. Transmission electron microscopy of KP-1S, KP-1S△*act*, KP-3R and KP-3R△*ompK26*. One representative image from four images obtained from one section is shown. C. Biofilm formation of KP-1S, KP-1S△*act*, KP-3R and KP-3R△*ompK26* in polystyrene plates. P values were calculated from student t-test.

Interestingly, the mucoid phenotype of KP-1S△*act* has reduced compared with KP-1S (**Figure S4**). Given that mucoid phenotype has been associated with the production of capsule polysaccharide, the transmission electron microscopy assay was performed (**Figure 4B**). Results showed that capsule production of KP-1S△*act* has reduced compared with KP-1S, demonstrating that *act* plays an important role in the synthesis of CPS in ST11-CRKP. As the capsule production of clinical isolates was inversely related to biofilm formation (15), biofilm was analyzed. Results showed that KP-1S△*act* has significantly increased the biofilm productions compared with its wild type strain (**Figure 4C**), which probably facilitates its long-term carrage and persistence in host (16).

### Transcriptome analysis identified genes affected by *act*

RNA-seq was employed to determine the transcriptomes of KP-1S, KP-3R, KP-1S△*act* and KP-3R△*ompK26* (**Table S3**). The heatmap showed that the transcriptome profiles of all the isolates were consistent within each group and most of the genes expressed uniformly across all the samples (**Figure 5A**). As shown in **Figure 5B**, the first principal component (PC1) and the second principal component (PC2) explained up to 91% of the variance of gene expression, indicating that only few genes were differentially expressed and contributed to phenotype changes.

**Figure 5.**
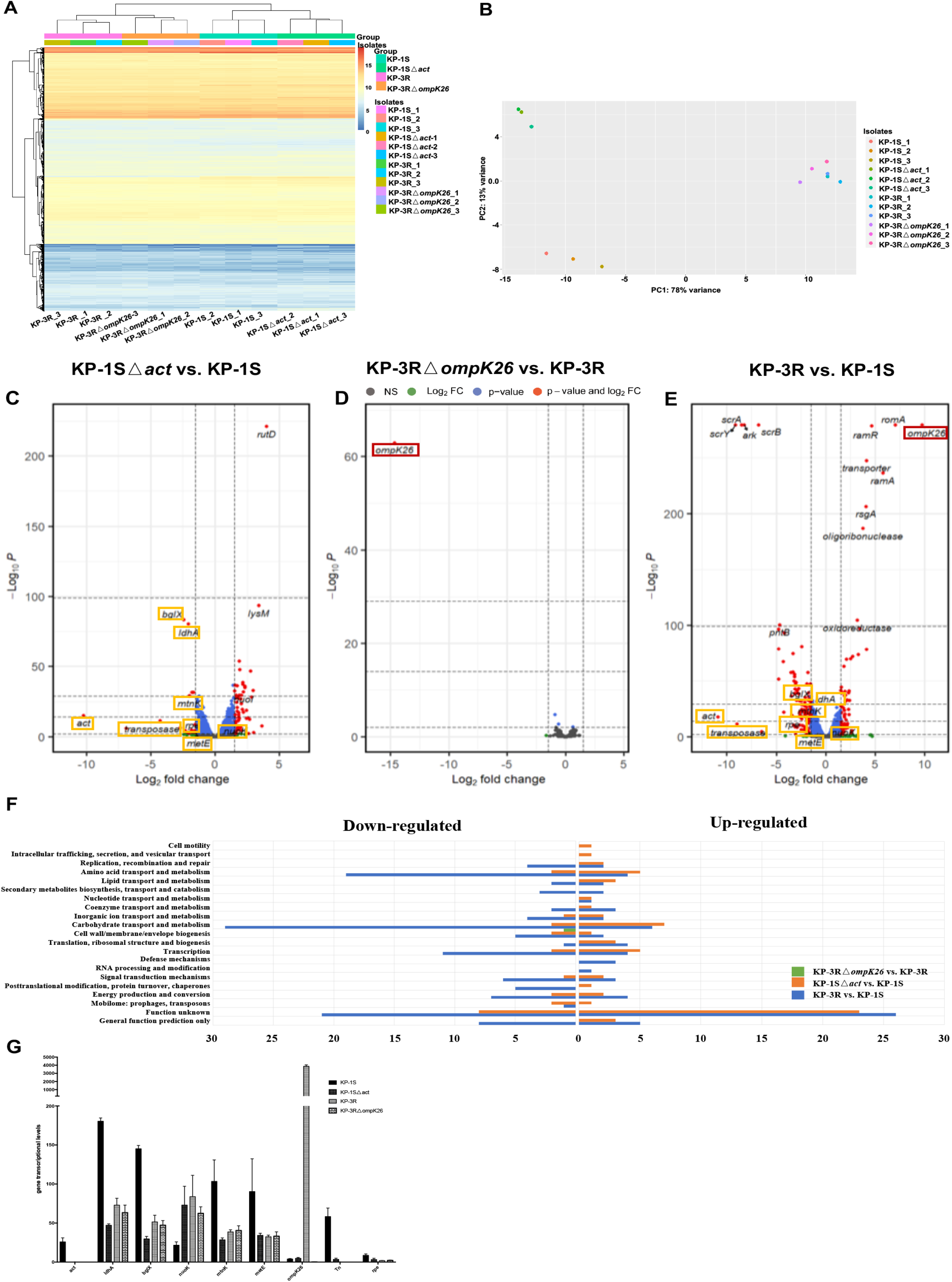
RNA-seq analysis of KP-1S, KP-3R, KP-1S△*act* and KP-3R△*ompK26.* A. The heatmap of transcriptional profiles of KP-1S, KP-3R, KP-1S△*act* and KP-3R△*ompK26.* B. Principal component analysis (PCA) of transcriptional profiles of KP-1S, KP-3R, KP-1S△*act* and KP-3R△*ompK26.* C. The volcano plot of differentially expressed genes in KP-1S△*act* vs. KP-1S. D. The volcano plot of differentially expressed genes in KP-3R△*ompK26* vs. KP-3R. E. The volcano plot of differentially expressed genes in KP-3R vs. KP-1S. Genes highlighted with rectangle were shared with other groups. F. COG analysis of the differentially expressed genes from each group. Genes with unknown functions were omitted. G. The transcriptional levels of genes differentially expressed in both KP-1S△act vs. KP-1S and KP-3R vs. KP-1S.

The differentially expressed genes were identified between KP-1S△*act* and KP-1S, KP-3R△*ompK26* and KP-3R, as well as KP-3R and KP-1S (**Table S4-6**). In the group of KP-1S△*act* vs. KP-1S, 21 genes were under-expressed and 61 genes were over-expressed in KP-1S△*act*. Besides *act*, other genes including *ldhA*, *bglX*, *mtnK* and *metE* were differentially expressed (**Figure 5C**). In the group of KP-3R△*ompK26* vs. KP-3R, only *ompK26* was under-expressed in KP-3R△*ompK26*, indicating that *ompK26* is located at the end of the pathway (**Figure 5D**). In the group of KP-3R vs. KP-1S, 115 genes were under-expressed and 66 genes were over-expressed in KP-3R (**Figure 5E**). Besides *act* and *ompK26*, *ldhA*, *bglX*, *mtnK* and *metE,* which were observed in KP-1S△*act* vs. KP-1S, were also differentially expressed.

COG functional analysis revealed that ‘cell wall/membrane/envelope biogenesis’ was affected in KP-3R△*ompK26* vs. KP-3R (**Figure 5F**). The top three affected functions in KP-1S△*act* are ‘carbohydrate transport and metabolism’, ‘amino acid transport and metabolism’, and ‘transcription’ compared with KP-1S. Interestingly, though with additional genomic differences in KP-3R, the top three affected functions in KP-3R vs. KP-1S are the same as those in KP-1S△*act* vs. KP-1S.

Given that differentially expressed genes presented simultaneously in KP-1S△*act* vs. KP-1S and KP-3R vs. KP-1S are more likely to be associated with the virulence phenotype, therefore, genes up-or down-regulated in KP-1S△*act* and KP-3R compared with KP-1S were identified. Besides *act*, genes including *ldhA*, *bglX*, *mtnK*, *metE*, transposon and *rpe* were down-regulated, and *nuoK* was up-regulated in KP-1S△*act* and KP-3R (**Figure 5G**). *ldhA* encodes for D-Lactate dehydrogenase A and participates in fermentative lactate dehydrogenation. *bglX* encodes for beta-glucosidase which hydrolyzes beta-D-glucosyl residues to beta-D-glucose. *mtnK* encodes for S-methyl-5-thioribose kinase and *metE* encodes for homocysteine S-methyltransferase. Both of them participated in the methionine synthase and methylation. *rpe* encodes for ribulose-phosphate 3-epimerase catalyzes the reversible epimerization of D-ribulose 5-phosphate to D-xylulose 5-phosphate, which is important for carbohydrate degradation. *nuoK* (also known as ND4L) encodes for NADH-quinone oxidoreductase subunit K and shuttles electrons from NADH to quinones in the respiratory chain. The results also showed that KP-3R yielded abundant transcripts of *ompK26*, suggesting that KdgR has a strong repressive effect on *ompK26*.

## DISCUSSION

In this study of ST11-CRKP sequentially isolated from a patient with scrotal abscess, the within-host genomic dynamics were deciphered. The study begins with ST11-CRKP strain that was initially susceptible then resistant to tigecycline during tigecycline therapy. Tigecycline resistant ST11-CRKP infections are generally considered fatal in clinical for their association with high mortality and poor outcomes (17). However, in this study, the patient dramatically recovered from the fatal infection without effective medication and ST11-CRKP strains can be continuely isolated from the urine of the patient. We found that the virulence of tigecycline resistant ST11-CRKP was attenuated compared to the initial tigecycline susceptible strain in the mice infection model, which restated the fitness cost of acquiring antibiotic resistance in *K. pneumoniae*. The estimated instantaneous mutation rate of ST11-CRKP in this study was 2.1×10^−6^-1.7×10^−5^ substitutions per site per year, which is higher than the reported 6.9×10^−7^-1.8×10^−6^ substitutions per site per year (18, 19). The mutation rate might be overestimated in this study due to the limited sampling timepoints and the continuously selective pressure from antibiotics therapy.

The mutations selected *in vivo* may have crucial impacts on disease outcome, therefore we analyzed genomic changes in ST11-CRKP strains to decipher the genetic basis for within-host adaptation. Studies of pathogen adaptation during infection focused predominantly on mutations within coding regions, whereas adaptive mutations in intergenic regions received less attention (20, 21). However, almost all the mutations identified in this study are located in the intergenic or regulatory regions and the functions of two intergenic mutations have been verified, which underlines the importance of intergenic mutations in within-host adaptation. A study of intergenic evolution of *Pseudomonas aeruginosa* revealed that intergenic mutations represent an important aspect of bacterial evolution in niche adaptation (22). Therefore, we considered that intergenic evolution might be a more cost-effective way than coding region evolution in the acquisition of novel phenotypes and mediating host adaptation.

We further analyzed the genomic mutations associated with phenotypic variations. The tigecycline resistance owing to deletion of *ramR* RBS has been discussed in our previous publication, here our focus is on two mutations, including the 5 bp (TGTTT)-deletion found on upstream of *ompK26* and the deletion containing *act*, which are likely related to virulence attenuation. We proved that the TGTTT deletion is located on the binding site of *ompK26* repressor KdgR, and the deletion increases the expression level of *ompK26* to a large extend. *ompK26* encodes for a KdgM family porin and a previous study showed that the knockout of *ompK26* increases the virulence and carbapenem resistance in *K. pneumoniae* (23). We showed that the knockout of *ompK26* slightly increases the virulence of CRKP but the Mantel-Cox log rank test showed no significance between the survival rates of KP-3R△*ompK26* and KP-3R infected mices. RNA-seq results showed that *ompK26* is located at the end of the pathway and no differentially expressed gene was found between KP-3R△*ompK26* and KP-3R besides *ompK26*.

*act* is located in the region of CPS synthesis gene cluster and was predicted to encode an acyltransferase family protein. CPS is present on the surface of both Gram-positive and Gram-negative bacteria and it is an important virulence factor mediating host immune response (24, 25). The acetylation of CPS is frequent (26), and studies have shown that CPS acetylation enhances antigenicity and increases immunogenicity in *Escherichia coli* (27), *Streptococcus agalactiae* (28) and *Neisseria meningitidis* (29). Besides, the acetylation of CPS increases mucoid colony and reduces the aptitude to biofilm formation in *E. coli* (30), which have been observed in the *act* deletion and knockout strains in our study. Previous study showed that deficiency in CPS biosynthesis increases biofilm formation, which makes the pathogens difficult to eradicate in urinary tract and facilitates dissemination (16). Research on CPS acetylation in *K. pneumoniae* type K57 showed that the acetylation enhanced the immunoreactivity of CPS and increased the induction of pro-inflammatory cytokines (31). The ST11-CRKP strains in this study belong to KL64, which was considered a virulence enhanced clone (6). Previous study of an *act*-harboring KL64 strain NCTC 9184 showed that a D-glucose of CPS was acetylated (32), indicating that CPS acetylation happens in KL64 *K. pneumoniae* strains. Given the fact that *act* deletion was found in the virulence attenuated isolates and knockout of *act* reduced the virulence in mutants, we inferred that *act* likely mediated the virulence variation by acetylating the CPS in this study.

Genes participated in lactate dehydrogenation (*ldhA*), beta-D-glucose synthesis (*bglX*) and methylation (*mtnK* and *metE*) were found under-expressed in *act*-abolished strains. The deletion of *ldhA* in *N. meningitidis* promotes biofilm formation (33), therefore, the increased biofilm formation in *act*-abolished strains might be mediated by the low expression of *ldhA*. As D-glucose synthesis and methylation are required for CPS biosynthesis, the decreased CPS productions in *act*-abolished strains were likely caused by the low expressions of *bglX*, *mtnK* and *metE*. Besides, the functions of other differentially expressed genes, such as *rpe* and *nuoK,* in *K. pneumoniae* were undetermined and will be further explored in the future.

In summary, our work illustrated the within-host evolution of a worldwide-disseminated clone ST11-CRKP from a clinical case by leveraging the power of WGS and building a direct connection between the genomic variants and host adaptation by RNA-seq and molecular biology techniques. Our results provide a better understanding of the evolutionary capacity and within-host adaptation of bacteria, which will be necessary for pathogens surveillance and infection-control in the future.

## MATERIALS AND METHODS

### Strains and growth conditions

KP-1S, KP-2S, KP-3R and KP-4R were isolated from the urine of a patient with scrotal abscess and urinary tract infection during antibiotics treatment (10). Other strains used in this study were constructed from KP-1S and KP-3R. All strains were cultivated in lysogeny broth (LB) medium at 37 °C, information of strains as indicated in **Table S1**.

### Mouse model of intraperitoneal infection

Male ICR mice (6-8 weeks old, weighing 20-25g) were infected intraperitoneally with 10^6^ CFUs of *K. pneumoniae* (10 mice/group) being harvested from the exponential growth phase. Mice were monitored for 7 days and assessed for death every 16-24 h. All animal experiments were performed following the protocols approved by the Animal Ethics Committee of Shanghai Skin Diseases Hospital.

### Whole genome sequencing and bioinformatics analysis

Genome sequencing was performed as described previously (11). Briefly, genomic DNA of KP-1S, KP-2S, KP-3R and KP-4R were extracted using bacterial genomic DNA extraction kit and sequenced using Illumina HiSeq 150-bp paired-end sequencing technologies. The sequencing reads were assembled using SPAdes V3.8 (34) with default parameters and contigs with less than 500 nucleotides were excluded. The genes were predicted and annotated using NCBI online annotation service. Genome alignments were performed by MUMmer 4 (35). The SNPs and INDELs were identified by Snippy v4.6 and CNOGpro with KP-1S as the reference. The phylogenetic relationship was constructed by the FastTree (36) based on the variants with the maximum-parsimony method. The genomic features including sequence typing, virulence genes, antimicrobial resistance genes, MLST and capsular type were analyzed by Kleborate v2.0.1. Plasmid replicons were identified by PlasmidFinder.

### Quantitative RT-PCR (qRT-PCR)

qRT-PCR was performed as described previously (10). Briefly, RNA were extracted from mid-log-phase bacterial cultures using the RNeasy mini kit (Qiagen). cDNA was synthesized using the RT reagent kit with gDNA eraser (Takara). qRT-PCR was performed using SYBR Premix ExTaq (TaKaRa) on a CFX96 Real-Time PCR Detection System (Bio-Rad). PCR primers for *ompK26* and the endogenous reference gene *rrsE* were provided in **Table S2**.

### Construction of *ompK26* and *act* mutant

Knockout of chromosomal *ompK26* and *act* was conducted as we described previously (37). Briefly, pKOBEG was transformed into KP-3R to generate Kp-3R/pKOBEG. Homology fragments of *ompK26* were amplified and inserted into the pMD-18T-hph on either side of the hygromycin gene. The recombinant plasmid was then digested by KpnI and HindIII to get the final linear fragment. The final fragment was transformed into Kp-3R/pKOBEG. Mutant clones were screened by PCR using the primer pairs of internal-F/internal-R and external-F/external-R. The same method was applied to construct *act* mutant. Strategy for constructing and identification of mutant clone are shown in **Figure S3**.

### Complementation of *ompK26* with its native promoter and *kdgR* overexpression

The *ompK26* and the native promoter region of KP-1S and KP-3R were amplified by *ompK26*_promoter_F/R (**Table S2**) and cloned into pMD-18T-hph at HindIII and KpnI sites to generate pMY53 and pMY54, respectively. pMY53 and pMY54 were electrically transformed into KP-3R△*ompK26* to generate KP-3R△*ompK26*/pMY53 and KP-3R△*ompK26*/pMY54. Strategies for construction pMY53 and pMY54 are shown in **Figure S2**. The full-length *kdgR* was amplified from KP-1S using pBAD33_*kdgR*_F/R (**Table S2**) and cloned into pBAD33 to generate pMY59. pMY59 was then transformed into KP-3R△*ompK26*/pMY53 and KP-3R△*ompK26*/pMY54. KP-3R△*ompK26*/pMY53/pMY59 and KP-3R△*ompK26*/pMY54/pMY59 were confirmed by qRT-PCR.

### Purification of recombinant KdgR and electrophoretic mobility shift assay (EMSA)

The full-length *kdgR* was amplified from KP-1S using pET28a_*kdgR*_F/R (**Table S2**) and cloned into pET-28a to generate pMY55. The KdgR-6xHis fusion protein was expressed in BL21 (DE3) with 0.2 mM of Isopropyl β- d-1-thiogalactopyranoside at 18 °C. Protein purification was performed as previously described (10). Protein purity was confirmed by sodium dodecyl sulfate-polyacrylamide gel electrophoresis (SDS-PAGE) analysis. The promoter regions of *ompK26* were amplified using *ompK26*_prob_F/R (**Table S2**). The KdgR/DNA complexes were mixed, incubated, electrophoresis and imaged according to the procedure described previously (38).

### Biofilm formation and transmission electron microscopy (TEM)

Biofilm production was determined as described (39). Briefly, 1 ul of overnight culture was inoculated into 100 ul of fresh LB broth in each well of untreated 96-well polystyrene plates. After 24 h incubation at 37 °C, the wells were washed four times with water and 150 μl of 0.1% crystal violet was added. After 10 min incubation, crystal violet was removed and the wells were washed six times with water. Then, 200 μl of 80% ethanol was added and the plate was incubated for 10 min at room temperature before determining the OD595 with a microplate reader. Transmission electron microscopy was performed by the Electron Microscopy Facility of Servicebio (Wuhan, China), and images were captured by HITACHI HT7800/HT7700.

### RNA sequencing and differential expression analysis

Total RNA was used as input material for the RNA sample preparations. RNA sequencing libraries were prepared according to the manufacturer’s protocol and sequenced on Illumina Novaseq platform. The genome and gene model annotation of KP-1S was used as the reference. The reads were mapped to the reference genome by Bowtie2 v2.4.2 (40). Differential expression analysis of two conditions/groups (three biological replicates per condition) was performed using DESeq (1.18.0) (41). P-values were adjusted using Benjamini and Hochberg’s approach. Genes with adjusted P-value <0.001 and |log2FoldChange| >1.5 were classified as differentially expressed. Clusters of Orthologous Groups of proteins (COGs) database was used to classify the differentially expressed genes.

### Statistical analysis

Statistical analysis was performed by R. The Mantel-Cox log rank test was used to compare the Kaplan-Meier survival curves and calculate the P values. Student T test was used to compare the biofilm productions and calculate the P values.

## Data availability

The sequences of KP-1S have been deposited in the DDBJ/ENA/GenBank under the bioproject PRJNA590579.

## ACKNOWLEDGMENTS

This work was supported by the National Natural Science Foundation of China [grant number 82072322 and 81572039], the Basic Research Project of Shanghai Science and Technology Commission [grant number 15JC1403000], Central Government Guided Science and Technology Development Project [YDZX20193100002868], the Shanghai Natural Science Foundation [grant number 19ZR1442800, 15ZR1437000, 16411961300, 17DZ2293300], Three-Year Initiative Plan for Strengthening Public Health System Construction in Shanghai [grant number GWV-10.2-YQ02], Clinical Research Plan of SHDC [grant number 16CR1029B] and National megaproject on key infectious diseases [grant number 2017ZX10202102-001–007].

M.Y., J.J. and P. Z. designed the study. M.Y. and J.J. drafted the manuscript. M.Y., C.L., M.S., D.Z., J.Y. and Z. H. carried out experiments. J.J. conducted the bioinformatic analyses. P.Z. and X.X. raised several useful suggestions. All authors read and approved the final manuscript.

We declare that we have no conflicts of interests.

